# NAD^+^ depletion and altered mitochondrial function are key to the establishment of placental dysfunction in an inflammatory-driven subclass of preeclampsia

**DOI:** 10.1101/2023.09.09.556974

**Authors:** Fahmida Jahan, Goutham Vasam, Yusmaris Cariaco, Abolfazl Nik-Akhtar, Alex Green, Keir J. Menzies, Shannon A. Bainbridge

**Affiliations:** Department of Biochemistry, Microbiology and Immunology, Faculty of Medicine, University of Ottawa, Ottawa, ON, Canada; Interdisciplinary School of Health Sciences, Faculty of Health Sciences, University of Ottawa,Ottawa, ON, Canada; Ottawa Institute of Systems Biology, University of Ottawa, Ottawa, ON, Canada; Department of Cellular and Molecular Medicine, Faculty of Medicine, University of Ottawa, Ottawa, ON, Canada

## Abstract

Preeclampsia (PE) is a pregnancy associated hypertensive disease. It is one of the major causes of pregnancy-related maternal/perinatal adverse health outcomes, with a lack of highly effective preventative strategies and/or therapeutic interventions. Our group has previously identified distinct subclasses of pathophysiology underlying a PE diagnosis, one of which exhibits heightened immune activation at the gestational parent-fetal interface, identified as inflammatory-driven PE. In non-pregnant populations, chronic inflammation is associated with reduced cellular availability of NAD^+^, a vitamin B3-derived metabolite involved in energy metabolism and mitochondrial function. Interestingly, specifically in placentas from women with inflammatory-driven PE, we observed increased activity of NAD^+^-consuming PARP enzymes and reduced NAD^+^ content. Moreover, these placentas had decreased expression of several mitochondrial oxidative phosphorylation (OXPHOS) proteins and evidence of oxidative damage. This human data was supported by cell culture findings, which likewise demonstrated increased PARP activity, coupled to decreased mitochondrial respiration rates and decreased invasive function of cultured HTR8 human trophoblast cells, following inflammatory induction by TNF-α. Importantly, these adverse inflammatory effects were attenuated by boosting cellular NAD^+^ levels with nicotinamide riboside (NR). Finally, using an LPS-induced rodent model of inflammatory-driven PE, we demonstrated that NR administration (200mg/kg/day) from gestational day (GD) 1-19 could prevent the development of maternal hypertension and fetal/placental growth restriction, improve placental mitochondrial function, reduce placental inflammation and oxidative stress. Thus, this study demonstrates the critical role of NAD^+^ metabolism in maintaining healthy placental function and identifies NAD^+^ boosting as a promising preventative strategy for the inflammatory-driven subclass of PE.

**One sentence summary:** Boosting NAD^+^ levels prevent inflammatory-driven preeclampsia by improving placental mitochondrial function.

## Introduction

Preeclampsia (PE) is a complex hypertensive disorder that occurs during pregnancy (at ≥20 weeks of gestation) [1]. The disease affects almost 10 million women each year, making it one of the major causes of mortality and morbidity in the mother and baby [1]. Unfortunately, there is no cure for PE. Removal of the affected organ, the placenta, is the only ‘cure’, often resulting in premature delivery and associated iatrogenic outcomes [2]. Historically, PE has been studied as a singular disease. However, there is a considerable heterogeneity in the PE patient population based on the clinical profile, onset of disease, pregnancy outcome, fetal growth impacts and placental histopathology. It is now acknowledged that PE likely develops because of several distinct underlying etiologies [3–7]. Our group has previously characterized three different subclasses of PE pathophysiology using a multiscale profiling approach that incorporated gestational-parent/fetal clinical features and detailed placental profiling (molecular and histological). Distinct subclasses of PE disease identified included: 1) Gestational parent-driven PE (PE1) – likely the result of underlying gestational parent predisposing factors (involving minimal placental pathology); 2) Hypoxia-driven PE (PE2) – resulting from placental ischemia-reperfusion injury (~ 60% of the PE patient population); and 3) Inflammation-driven PE (PE3) – resulting from aberrant inflammation at the gestational parent-fetal interface [3, 4]. Existence of such disease subclasses necessitates the requirement for studies focused on better understanding subclass-specific disease processes and/or subclass-specific interventions. To date, most research on PE has focused almost exclusively on understanding the hypoxia-driven pathophysiology of PE, while the etiological relevance of parental factors and/or inflammatory processes have been given less attention. The current study aims to address this gap in knowledge, specifically focusing on better understanding placental dysfunction associated with inflammation-driven PE, and subclass-specific therapeutic interventions for this unique patient population.

Placental mitochondrial dysfunction has long been recognized as a key component of PE pathophysiology [8–12]. Altered placental mitochondrial respiration and/or total content is observed in tissues collected from pregnancies that span the clinical spectrum of PE, associated with altered oxidative phosphorylation (OXPHOS) protein expression, mitochondrial structural damage, defects in fusion/fission dynamics and increased oxidative stress [8, 13–15]. There is strong evidence to suggest that placental mitochondrial dysfunction may be a common feature across all PE subclasses [8], however a lack of subclass-specific research endeavours has limited our current understanding of this pathology, specifically in the context of inflammation-driven PE. In non-pregnant populations, several inflammatory disease conditions have been tightly linked to mitochondrial dysfunction, in large part driven by profound overactivity of nicotinamide adenine dinucleotide (NAD^+^) consuming enzymes, leading to depletion of intracellular NAD^+^ stores [16–28]. In the current study, this body of work will be expanded to determine if NAD^+^ depletion may likewise initiate placental mitochondrial dysfunction in inflammation-mediated PE.

NAD^+^, a vitamin B3 redox metabolite, is a critical coenzyme central to metabolic function throughout the body [29–31]. NAD^+^ maintains mitochondrial health by regulating energy metabolism and by inducing pathways, such as the mitochondrial unfolded protein response [29–31]. Enzymes such as poly-adenosine diphosphate (ADP)-ribose polymerases (PARPs), Sirtuins (SIRT), cluster of differentiation 38/157 (CD38/CD157), and sterile alpha and toll/interleukin-1 receptor motif-containing protein 1 (SARM1), require NAD^+^ as a co-substrate to carry out their enzymatic functions [19, 29–31].

Among these, two of the most well-described NAD^+^-consuming enzyme families are the PARPs and the SIRTs. A primary function of PARPs is the post-translational modification of proteins, via the covalent addition of ADP-ribose polymers to various proteins – a process known as poly(ADP-ribosyl)ation (PARylation). This NAD^+^-dependant process helps to regulate biological processes such as chromatin organization, DNA damage repair, mRNA stability, transcriptional control, DNA methylation, glycolysis and inflammation [32–35]. SIRTs, on the other hand, are known to be vital for mitochondrial metabolism and biogenesis, and ROS detoxification [19, 29–31]. Under normal conditions, cellular NAD^+^ content is tightly regulated, achieving an optimal balance between NAD^+^ biosynthesis/salvage and NAD^+^ consumption pathways. However, under proinflammatory conditions some of the major NAD^+^ consumers, such as PARPs, become highly activated, tilting the scales of NAD^+^ homeostasis towards excessive consumption and quickly deplete intracellular NAD^+^ stores, leading to mitochondrial dysfunction and impaired energy metabolism [16–28, 30, 36].

Interestingly, in a mouse model of hypoxia-drive PE, replenishment of whole-body NAD^+^ stores via nicotinamide (NAM) treatment have been shown to rescue several PE-like features (i.e. hypertension, fetal growth restriction) [37, 38]. However, the role of placenta-specific NAD^+^ depletion in the pathogenesis of PE and whether supplementation with an NAD^+^ precursor will be an attractive preventative strategy for other PE subclasses is unknown.

In the current study, the potential role(s) of inflammation-mediated NAD^+^ depletion and subsequent mitochondrial dysfunction in the establishment of placental disease in PE were explored using complementary studies on human placenta tissues, a human trophoblast cell culture model and in a rat model of inflammation-driven PE. Further, the therapeutic potential of NAD^+^ boosting, via nicotinamide riboside (NR) treatment, on placental health, fetal growth and clinical manifestations of inflammation-mediated PE were evaluated.

## Results

### Human placenta tissues from inflammation-driven PE demonstrate evidence of PARP-mediated NAD^+^ depletion and mitochondrial dysfunction

Previous gene set enrichment carried out by our group using a genome-wide microarray dataset from human placenta samples (Table 1) annotated according to PE subclass, identified an overexpression of several pro-inflammatory signaling pathways uniquely within one PE subclass – labelled inflammation-mediated PE (PE3) [3, 4]. Using this same dataset, here we show that this pro-inflammatory gene expression profile is coupled to elevated expression of several *PARP* transcripts uniquely in these same PE3 cases, findings not observed in the placentas from the other PE subclasses (PE1 and PE2; Fig 1A). PARP hyperactivation has been widely described in non-pregnant inflammatory conditions, leading to excess protein PARylation and NAD^+^ depletion [16–28, 30, 36]. As such the degree of protein PARylation was assessed in human placentas collected from each of the three PE subclasses, along with term, preterm and chronic hypertension controls. Global protein PARylation was also found to be uniquely higher in the placentas from PE3, compared to other PE subclasses and all control groups (Fig 1B-C). In parallel, LC/MS measurements demonstrated significant depletion of NAD^+^ and total NAD(H) in PE3 placentas, compared to control preterm and term placentas (Fig 1D-E). No significant reduction was observed for NAD^+^ and total NAD(H) content in the other PE subclasses (PE1 and PE2) or in the chronic hypertension groups (Fig 1D-E). NADH levels alone, the NAD^+^/NADH ratio and NAM, an end product of NAD^+^ consumption, on the other hand, did not change in any of the groups (Fig 1F-G, Supplementary Fig 1A).

**Figure 1.**
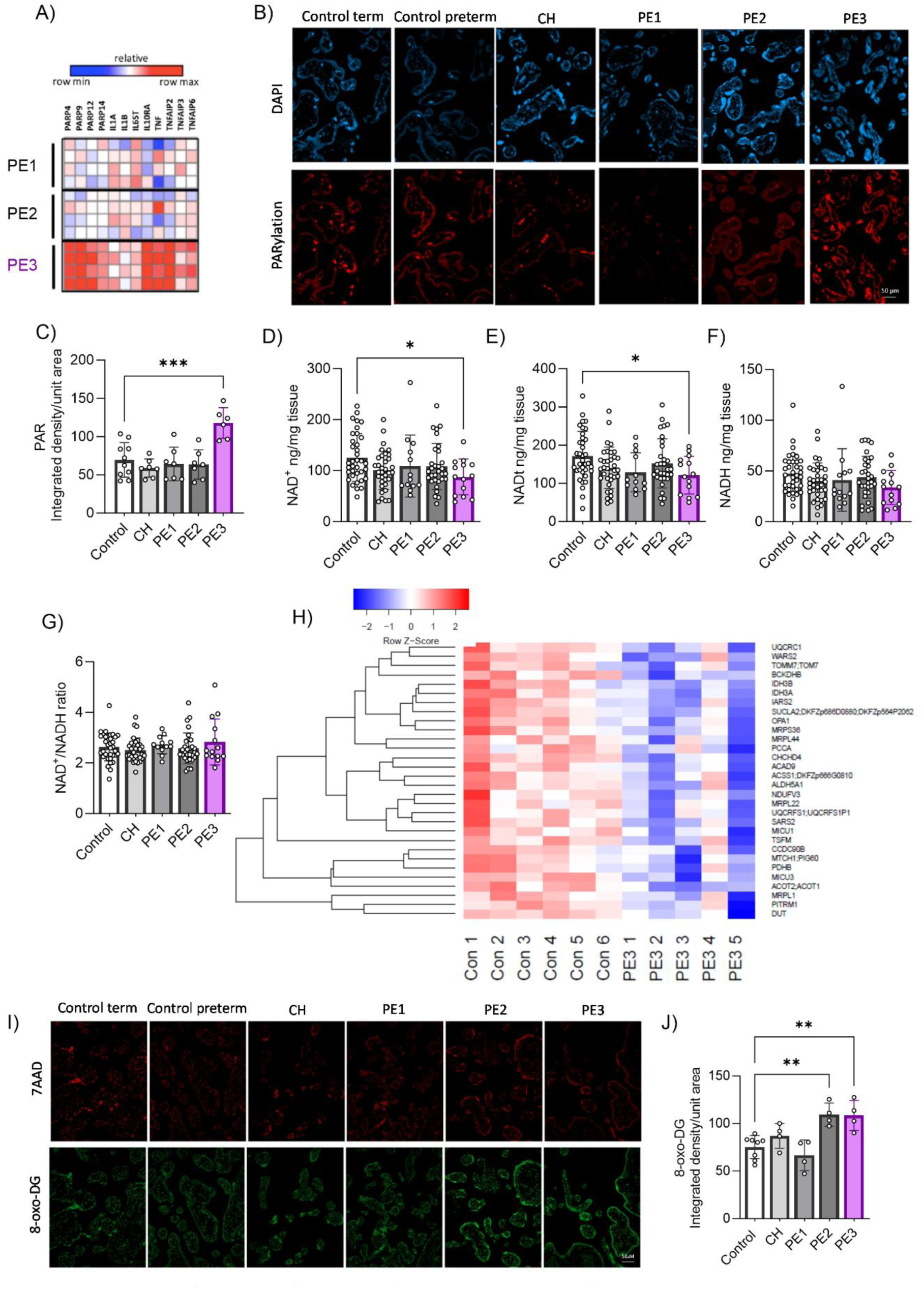
Identification of altered NAD^+^/PARP-signalling, mitochondrial proteomes and oxidative damage in human placentas with inflammation-driven PE (PE3). A) Human placenta gene expression of inflammatory markers and NAD^+^ consuming enzymes in human placenta biopsies from all three PE subclasses (n=4 placentas/group), B-C) Representative immunofluorescence images and quantification of protein PARylation in human placenta tissue sections from all three PE subclasses and various control groups (n=7-8 placentas/group). D-G) LC/MS quantification of human placental NAD(H) levels (n=14-36 placentas/group). H) Heatmap of downregulated mitochondrial proteins in inflammatory PE3 (term) placentas compared to control term placentas (n=5-6 per group). Shown are only proteins that met the False Discovery Rare (FDR) threshold of 0.05. I-J) Placenta oxidative stress was determined by immunofluorescence staining of human placenta tissue sections with anti-8 oxoDG antibody and quantification by integrated intensity/area of trophoblasts (n=4 per group). One-way ANOVA with Holm-Šídák’s multiple comparisons test *P<0.05 and **P<0.01. Error bar indicates standard deviation (SD). Control = control term and preterm, CH = chronic hypertension control, PE1 = Gestational parent-driven PE subclass, PE2 = Hypoxia-driven PE subclass, PE3 = Inflammation-driven PE subclass.

**Table 1:** Study population demographics.

|  | **Control**<br>**(preterm and**<br>**term)**<br>**(n = 43)** | **Chronic**<br>**hypertension**<br>**(n = 23)** | **PE1**<br>**(n = 16)** | **PE2**<br>**(n = 42)** | **PE3**<br>**(n = 17)** |
| --- | --- | --- | --- | --- | --- |
| **Maternal Age<sup>a</sup>** | 31.4±4.7 | 36.1±4.6<sup>c</sup> | 32.2±5.4 | 33.1±6.3 | 31.4±4.2 |
| **Gestational Age**<br>**at Delivery**<br>**(weeks)<sup>a</sup>** | 34.1±5.3 | 33.2±4 | 35±3.1 | 31±3.1<sup>c</sup> | 33.3±2.80 |
| **Fetal Sex (F)<sup>b</sup>** | 21/43 | 14/23 | 7/16 | 23/42 | 7/16 |
| **Birthweight (g)<sup>a</sup>** | 2481.3±1115.09 | 1649.8±867.7<sup>d</sup> | 2292.2±813.1 | 1290.6±611.1<sup>e</sup> | 1707.7±602.7<sup>c</sup> |
| **Birthweight %ile**<br>**(<10<sup>th</sup> %ile)<sup>b</sup>** | 2/43 | 15/23 | 5/16 | 30/42 | 10/17 |
| **Placental Weight**<br>**(g)<sup>a</sup>** | 541.09±180.4 | 344.1±164.8<sup>e</sup> | 454.7±133.5 | 312.3±128.2<sup>e</sup> | 372.7±147.1<sup>d</sup> |
| **Placenta**<br>**Hypoplasia**<br>**(<10<sup>th</sup> %ile)<sup>b</sup>** | 0/43 | 9/23 | 1/16 | 15/42 | 4/16 |
| **Maximum**<br>**Maternal Systolic**<br>**Blood Pressure<sup>a</sup>** | 118.4±15.4 | 162.8±19.6<sup>e</sup> | 158.3±15<sup>e</sup> | 162±16.9<sup>e</sup> | 150.7± 15.6<sup>e</sup> |
| **Maximum**<br>**Maternal**<br>**Diastolic Blood**<br>**Pressure<sup>a</sup>** | 71.9±11.6 | 101±11.6<sup>e</sup> | 100.7±7.8<sup>e</sup> | 103.7±12.8<sup>e</sup> | 95.8±11.3<sup>e</sup> |
| **Mode of Delivery**<br>**(C-section)<sup>b</sup>** | 21/43 | 20/23 | 9/16 | 40/42 | 16/17 |

<sup>a</sup> For mean ± standard deviation; <sup>b</sup> for number of pregnancies out of total number per group; <sup>c</sup> for p value <0.01; <sup>d</sup> for p value <0.001 and <sup>e</sup> for p value <0.000

Placental mitochondrial dysfunction and oxidative stress have been described as key contributors to placental dysfunction in PE when considered as a single clinical disease [8–12]. Thus, we aimed to determine if a reduced NAD^+^ pool in PE3 placentas may be associated with placenta mitochondrial dysfunction and/or oxidative stress. We found no indications of altered placental mitochondrial content between any of the three PE subclasses and all control groups when using citrate synthase expression and mitochondrial complex IV (COX-IV) activity assays as surrogates of mitochondrial content [39, 40] (Supplementary Fig 1B-C). However, using a proteomic approach on a subset placenta samples, numerous mitochondrial proteins were found to be down-regulated in PE3 placentas compared to controls, including: subunits of OXPHOS protein complexes [NADH:Ubiquinone Oxidoreductase Subunit V3 (NDUFV3), Ubiquinol-Cytochrome C Reductase Core Protein 1 (UQCRC1), UQCR Rieske Iron-Sulfur Polypeptide 1(UQCRFS1)], and proteins that effect mitochondrial dynamics [Optic atrophy type 1 (OPA1)], mitochondrial import (TOMM7), mitochondrial translation (mitochondrial ribosomal proteins, MRPS36, MRPL44, MRPS36, MRPL22, MRPL1), and others (Fig 1H). Placenta samples from cases of hypoxia-drive PE (PE2) likewise demonstrated a decrease in some mitochondrial proteins compared to controls, such as OXPHOS protein-ATP synthase membrane subunit f (ATP5J2), mitochondrial translation elongation factor (TSFM), MRPL44 etc. (Supplementary Fig 1D). However, placentas from parental-driven PE (PE1) did not show any changes in mitochondrial protein profiles compared to healthy controls.

Mitochondrial impairment is thought to be linked to oxidative stress and placental dysfunction in PE [8–12], however this relationship has not been explicitly examined in a subclass-specific manner. For this purpose, we performed 8-Oxo-2’-deoxyguanosine (8-oxo-DG) staining - detecting oxidized guanine residues in DNA [41] - of placenta sections collected from all three PE subclasses. Interestingly, both hypoxia-driven PE (PE2) and inflammation-driven PE (PE3) exhibited increased levels of oxidized nucleotides, when compared to parental-driven PE (PE1) and controls (Fig 1I-J).

### Boosting NAD^+^ decreases protein PARylation and prevents human trophoblast cell dysfunction under inflammatory culture conditions

To examine if inflammation-mediated NAD^+^ depletion alters human trophoblast health and function, a TNF-α induced inflammatory insult was applied to the SVneo/HTR8 trophoblast cell culture model – representative of the invasive extravillous cytotrophoblast population, long considered a key trophoblast population contributing to the establishment of PE pathophysiology [42–45]. Following 24 hrs of treatment with 10ng/ml of TNF-α, we observed a significant increase in protein PARylation (Fig 2A-B), coupled to a decrease in total cellular NAD^+^ levels and NAD^+^/NADH ratio (Fig 2C).

**Figure 2.**
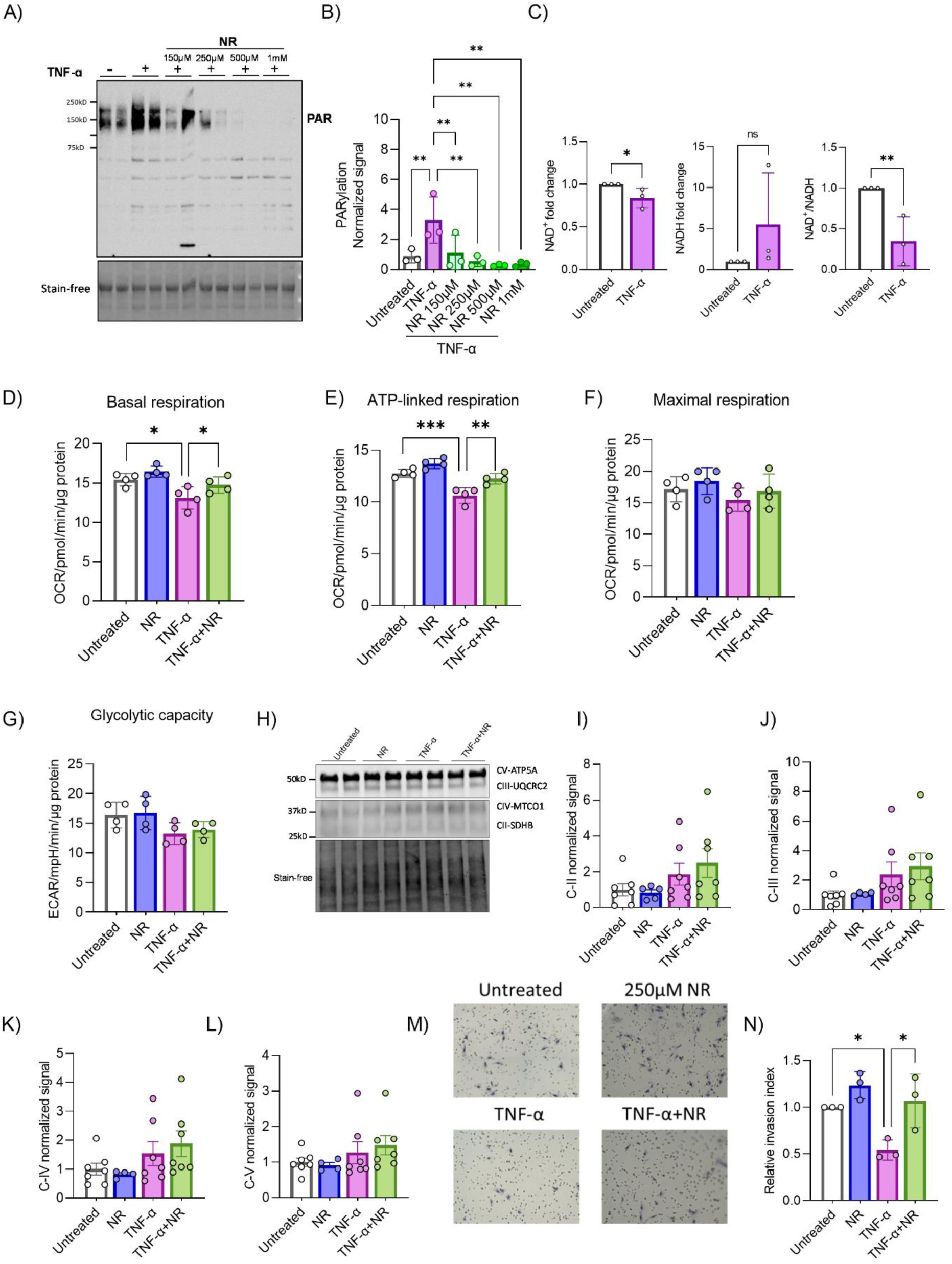
Potential of NAD^+^ boosting to improve HTR8/SVneo human trophoblasts health and function under an inflammatory *in vitro* condition. A-B) Trophoblast protein PARylation was determined by western blot with anti-PAR antibody on cells treated with 10ng/ml TNF-α and/or 150μM-1mM NR for 24 hrs (n=3). One-way ANOVA with Holm-Šídák’s multiple comparisons test C) Total cellular NAD^+^(H) and NAD^+^/NADH ratio quantification was performed on cells treated with or without 10ng/ml TNF-α for 24 hrs (n=3). One-tailed unpaired t-test. D-G) XFe96 seahorse Mito stress test was performed on cells treated with 10ng/ml TNF-α and/or 250μM NR for 24 hrs. Basal, ATP-linked and maximal mitochondrial respiration rates and extracellular acidification rates were measured (n=4). H-L) Expression of oxidative phosphorylation (OXPHOS) proteins were determined by western blot with OXPHOS antibody cocktail on cells treated with 10ng/ml TNF-α and/or 250μM NR for 24 hrs (n=4-7). M-N) Trophoblast invasion capacity was determined by performing Matrigel invasion assay on cells treated with 10ng/ml TNF-α and/or 250μM NR for 72 hrs (n=3). Two-way ANOVA with Holm-Šídák’s multiple comparisons test. *P<0.05,**P<0.01,***P<0.001. Error bar indicates standard deviation (SD). TNF-α= Tumor necrosis factor, NR= nicotinamide riboside.

Interestingly, when intracellular NAD^+^ content was boosted via NR treatment (150uM-1mM; Supplementary Fig 2), despite the continued presence of an inflammatory insult (TNF-α treatment), excessive protein PARylation was attenuated (Fig 2A-B). This would indicate that higher intracellular NAD^+^ concentrations are either inhibiting inflammation-mediated protein PARylation or are promoting faster flux of protein (de)PARylation. To determine if inflammation-mediated NAD^+^ depletion had any effect on mitochondrial function in this model, an XFe96 seahorse assay was conducted to measure mitochondrial oxygen consumption rates [46]. TNF-α treatment caused a decrease in basal and ATP-linked respiration, while NAD^+^ boosting with NR co-treatment was able to rescue these respiration rates (Fig 2D-E). No difference in maximal respiratory capacity was observed with either TNF-α or NR co-treatment (Fig 2F). Simultaneously, glycolytic capacity of these cells was assessed via measurements of extracellular acidification rates. However, there was no significant decrease in glycolytic capacity in any treatment groups (Fig 2G). An assessment of the expression of key OXPHOS subunit proteins [succinate dehydrogenase B (SDHB), Ubiquinol-Cytochrome C Reductase Core Protein 2 (UQCRC2), Mitochondrially Encoded Cytochrome C Oxidase I (MTCO1), ATP synthase subunit alpha of Complex V (vATP5A)], revealed no change in protein content with NR co-treatment, suggesting the improved mitochondrial respiration rates observed are likely related to mitochondrial functional capacity improvements rather than mitochondrial content improvements (Fig 2H-L). Finally, an assessment of the impact of inflammation-mediated NAD^+^ depletion on trophoblast function was carried out using a matrigel-coated boyden chamber invasion assay. The primary function of the extravillous cytotrophoblast cell lineage is invasion of the decidua and remodelling of the uterine spiral arterioles, ensuring adequate utero-placental blood supply to support fetal growth [47]. Previous studies have demonstrated an inhibitory effect of TNF-α treatment on the invasive capacity of these cells using either the HTR8 cell culture model or first trimester *ex vivo* placenta explants [48–50].

We were able to replicate these findings, demonstrating a 50% reduction in trophoblast invasion with TNF-α treatment. Importantly, intracellular NAD^+^ boosting with NR co-treatment rescued this deficit, returning the invasive capacity of these inflammatory insult-exposed cells back to those observed in untreated controls (Fig 2M-N).

Collectively, these cell culture findings providing preliminary evidence which suggests that NAD^+^ replenishment via NR treatment may improve human trophoblast health and function under pro-inflammatory conditions.

### Boosting NAD^+^ levels during pregnancy using an NR therapeutic intervention prevents the development of PE-like features in rat model of inflammation-driven PE

As NAD^+^ depletion was observed in placentas from cases of inflammatory-driven PE, and NAD^+^ supplementation *in vitro* improved human trophoblast health and function under pro-inflammatory conditions; we next sought to determine if NR could be used therapeutically to prevent the development of PE-like clinical features in a rat model of inflammation-driven PE. Our group has previously characterized and compared several rat models of inflammation-mediated PE, determining the lipopolysaccharide (LPS)-induced rat model was able to most closely recapitulating features observed in cases of human inflammation-driven PE [51]. In this model, LPS (20-70ug/kg/day) is injected daily from gestational day (GD) 13-18 intraperitoneally (IP). In our therapeutic intervention group, NR was administered using a preventative strategy by oral gavage from GD 1-19 (200mg/kg/day). Control pregnant rats received oral gavage of drinking water and saline IP injections using the same schedule (Fig 3A). As previously described, maternal BP was heightened at the end of pregnancy (GD 17-19) with LPS treatment, compared to saline controls, however this phenotype was rescued with NR intervention (Fig 3B).

**Figure 3.**
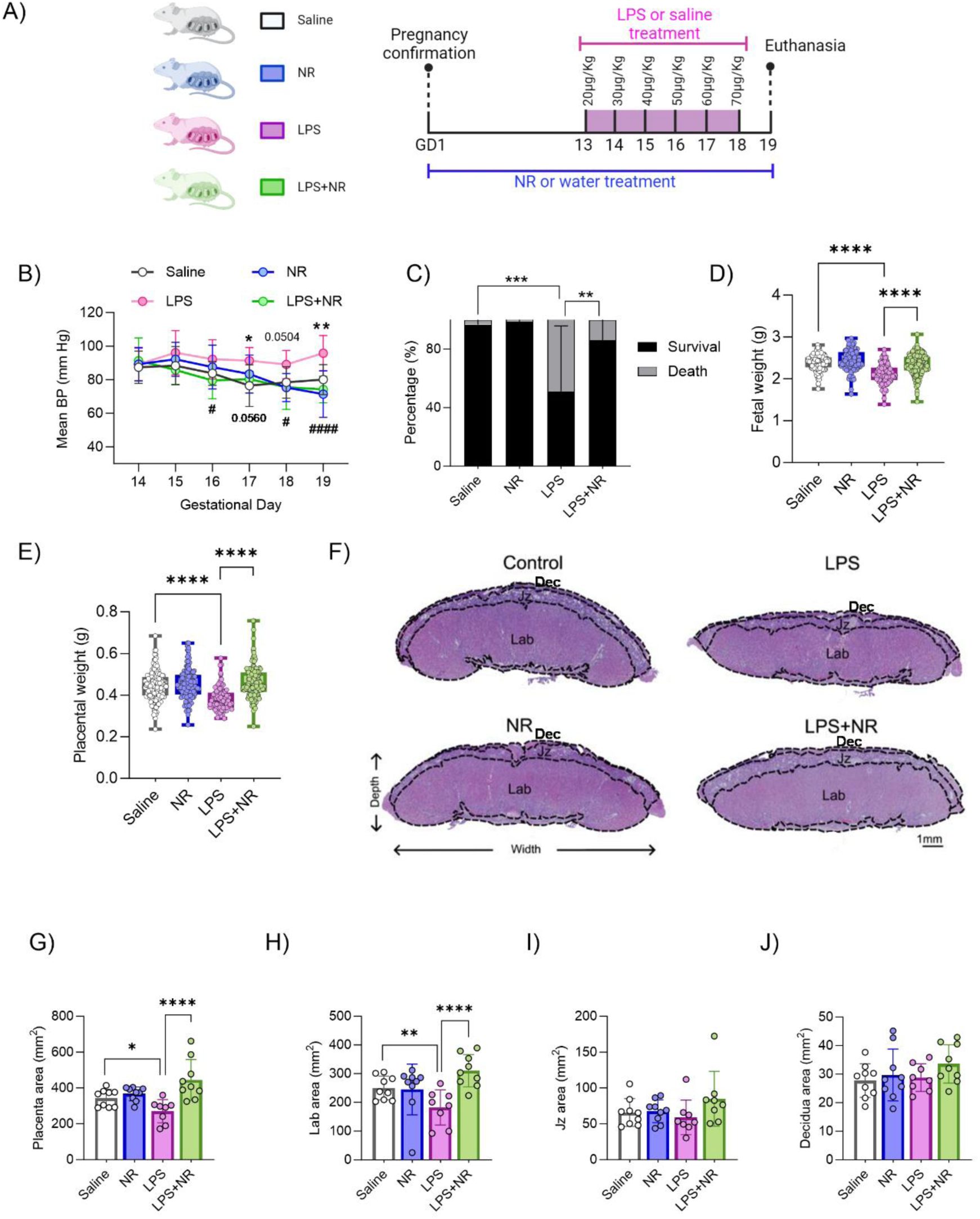
Effect of NAD^+^ boosting in preventing inflammation-driven PE in a rat model. A) Study design: Pregnant rats were given either NR or drinking water from GD1-19 and either LPS (20-70ug/kg/day) or saline were injected from GD13-18 and euthanasia was performed on GD 19 (Created with BioRender.com on June 12^th^, 2023). B) Blood pressure was measured from GD 14-19 by tail-cuff method (n=6-8 pregnant rats/treatment). * Indicates P values between saline and LPS and # indicates p value between LPS and LPS+NR. C) Fetal survival and death percentage. Two-way ANOVA with Holm-Šídák’s multiple comparisons test. D-E) Fetal and placental weights were measured (n=10 litters/treatment). One-way ANOVA with Holm-Šídák’s multiple comparisons test. F-J) Total placenta area along with individual regions such as labyrinth (Lab), junctional zone (Jz) and decidua were measured (n=9-10 litter/treatment, 1 placenta/litter). Two-way ANOVA with Holm-Šídák’s multiple comparisons test, */^#^P<0.05,**P<0.01,***P<0.001 and ****/^####^P<0.0001. Error bar indicates standard deviation (SD). LPS= Lipopolysaccharide, NR= nicotinamide riboside.

In the LPS-treated pregnancies, there was a significant decline in fetal survival to term (50.88%, Fig 3C), with the remaining viable fetuses demonstrating evidence of fetal growth restriction (Fig 3D). Importantly, these adverse fetal outcomes were attenuated with NR intervention (Fig 3C-D). Placental weights were decreased with LPS treatment (Fig 3E), with morphometric analysis demonstrating a decrease in total placenta area (Fig 3F-G), largely attributed to decreased size of labyrinth compartment of the placenta (Fig 3H) – the site of maternal-fetal exchange. NR intervention during pregnancy was therefore able to rescue this adverse LPS-induced placenta phenotype. On the other hand, the area of junctional zone (the hormone production area of the placenta) and the maternal decidua did not change with any treatment group (Fig 3I-J).

### Boosting placenta NAD^+^ levels using an NR therapeutic intervention prevents PARP-mediated NAD^+^ depletion and mitochondrial dysfunction in a rat model of inflammation-driven PE

To determine whether LPS treatment can induce placental inflammation, we first examined placental TNF-α levels by ELISA. TNF-α is considered to be a key contributor to the pathogenesis of hypertension in pregnancy [52–54]; and according to our microarray data, the TNF-α pathway is upregulated in the inflammatory PE3 placentas (Fig 1A). Similarly, we show that LPS treatment induces rat placental TNF-α expression, which can be normalized with NR intervention (Fig 4A). RNA-seq analysis performed on a subset of placentas further confirm that NR treatment in the presence of LPS modulates the placenta transcriptome (Supplementary Fig 3A-D) and downregulates defense response and response to chemokine (Supplementary Fig 3D) suggesting a reduction in inflammation.

**Figure 4.**
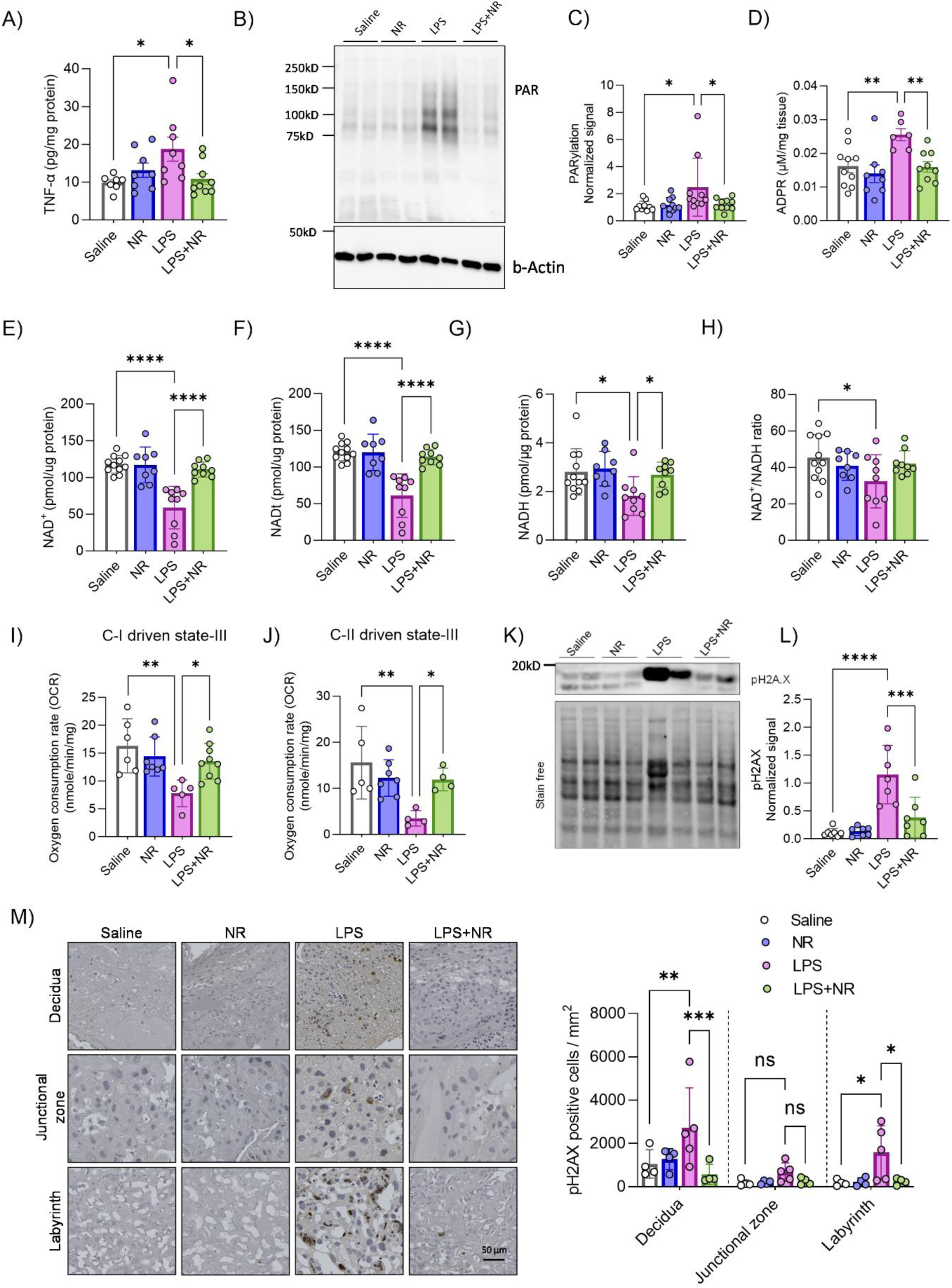
Effect of NAD^+^ boosting on placental dysfunction in a rat model of inflammatory PE. A) Placental TNF-α levels were determined by ELISA (n=7-9 litter/treatment, 1 placenta/litter). B-C) Placental protein PARylation was determined by western blot with anti-PAR antibody (n=10 litter/treatment, 1placenta/litter). D) Placental ADPR levels were determined by LC/MS (n=6-10 litter/treatment, 1placenta/litter). E-H) Placental NAD^+^(H) levels were determined by a colorimetric assay (n=8-10 litter/treatment, 1placenta/litter). I-J) Placental mitochondrial function was determined by performing Oxygraph assay on isolated mitochondria. Complex-I and II mediated oxygen consumption rates were recorded to determine their activity (n=4-8 litter/treatment, 6-8 placentas/litter). K-L) Placental oxidative stress was determined by western blot with anti-pH2A.X antibody, a marker of oxidative DNA damage (n=7 litter/treatment, 1placenta/litter). M-N) Placental sections were stained with anti-pH2A.X antibody to determine the number of pH2A.X positive cells in different regions of placenta (n=5 litter/treatment, 1placenta/litter). Two-way ANOVA with Holm-Šídák’s multiple comparisons test, *P<0.05,**P<0.01,***P<0.001. Error bar indicates standard deviation (SD). ADPR= ADP ribose, LPS= Lipopolysaccharide, NR= nicotinamide riboside.

To test if placental PARylation is hyperactivated in inflammatory PE, and if it causes a decline in NAD^+^ content, we examined rat placentas from LPS-induced PE for PARylated proteins and NAD^+^ content. We found that LPS led to increased placental protein PARylation while NR intervention normalized this change (Fig 4B-C). The levels of ADP-ribose (ADPR), a moiety produced from the hydrolysis of PAR chains by ADP-ribosyl-acceptor hydrolases [55, 56], were higher in placentas from LPS-induced rats and normalized with NR intervention, indicating that NR intervention likely did not increase PAR hydrolysis to result in lower PARylation levels (Fig 4D). We further found a decrease in NAD^+^, NADt (NAD^+^ and NADH), NADH and the NAD^+^/NADH ratio in placentas from LPS treated pregnant rats, compared to saline controls, and an NR intervention prevented such declines in NAD^+^, NADt (NAD^+^ and NADH) and NADH levels (Fig 4E-G). However, NAD^+^/NADH ratio was not significantly rescued with NR supplementation (Fig 4H). Interestingly, during inflammation an increase in the expression of the NAD^+^-dependent cADPR synthase CD38 produces ADPR and causes a reduction in NAD^+^ levels [57, 58]. We, however, did not observe any significant increase in CD38 expression in LPS treated placentas, suggesting that PARPs are likely the primary driver of placental NAD^+^ consumption with LPS treatment (Supplementary Fig 4A-B). Given that NR alone did not alter NAD(H) levels in rat placentas, we measured the levels of NAM, which is a by-product of NAD^+^ utilizing enzymatic reactions, which can be converted back to NAD^+^ by the salvage pathway [31]. As expected, we found high levels of NAM in both NR alone and NR with LPS treated placentas (Supplementary Fig 4C).

To determine if LPS treatment leads to placental mitochondrial dysfunction and whether NR exerts its beneficial effect in part by improving mitochondrial function, we performed oxygraphy respirometry on freshly isolated mitochondria from these placentas. LPS treatment decreased complex I and II driven state-III respiration rates while NR intervention normalized respiration rates (Fig 4I-J). Complex IV mediated respiration was unaltered by LPS or NR treatments (Supplementary Fig 4D). Similar to measurements in human HTR8 cells, these functional changes were not a result of altered expression of NADH:Ubiquinone Oxidoreductase Subunit B8 (NDUFB8), UQCRC2, vATP5A OXPHOS proteins (Supplementary Fig 4E-H). Given that placental mitochondrial fusion is impaired in PE [8], and that the expression of the fusion protein OPA1 was reduced in human inflammatory PE placentas (Fig 1H), we examined the expression of OPA1 in all conditions. Although OPA1 protein expression is significantly reduced in placentas from LPS treated rats, NR intervention did not attenuate this effect (Supplementary Fig 4I-J). A compromised mitochondrial quality control pathway has been shown to contribute to placental dysfunction in canonical/hypoxia driven PE2 [59]. Thus, we determined the expression levels of two mitochondrial quality control proteases-YME1 like 1 ATPase (YME1L1) and Caseinolytic Mitochondrial Matrix Peptidase Proteolytic Subunit (CLPP). Both YME1L1 and CLPP expression levels were unaltered in each of the different treatment conditions (Supplementary Fig 4K-M).

As chronic inflammation and mitochondrial dysfunction can result in oxidative stress [60–63] and our results show that human inflammatory PE placentas exhibit more oxidative DNA damage (Fig 1I-J), we next investigated if LPS treatment leads to rat placental oxidative DNA damage. We performed western blotting to determine the levels of the phosphorylated form of histone H2A.X (p-H2A.X), a marker of DNA double stranded break [64]. Placentas from LPS treated rats exhibit higher levels of phospho-H2A.X protein, while NR intervention significantly attenuated this effect (Fig 4K-L). We further confirmed this by immunohistochemistry showing that LPS treated placentas have more p-H2A.X positive cells in the labyrinth and maternal decidua, whereas NR intervention normalized this effect to control levels (Fig 4M-N). This is consistent with studies that have shown that NAD^+^ replenishment can lead to decreased p-H2A.X levels indicating a reduction in DNA damage [65].

## Discussion

We show for the first time that NAD^+^ levels are depleted in both human and rat placentas that are affected by inflammation. Our results indicate that this reduction in NAD^+^ may be precipitated by the hyperactivity of NAD^+^-consuming PARP enzymes. *In vitro*, we have showed that boosting NAD^+^ levels with NR can improve trophoblast health and function under inflammatory conditions. Using the LPS-induced inflammatory PE rat model, we demonstrated that an oral pre-treatment with NR prevents the development of PE-like clinical features, evidenced by lowered maternal blood pressure, decreased inflammation, increased placental and fetal weights; and increased fetal survival.

We have observed that increased protein PARylation is unique to human inflammation-driven PE (PE3) placentas, findings that were replicated in placentas from LPS treated pregnant rats. Currently we do not know which PARPs are important in the context of inflammatory PE. Among the 17 members of PARP family, PARP1/2/5a/5b are known to have PARylating activity, while rest of the PARP family (PARP3/4/6/7/8/10/11/12/14/15/16) are associated with mono-ADPribosylation (MARylation). In contrast, PARP9/13 have no enzymatic activity [33]. Among these, PARP1-a major NAD^+^ consumer of the cell, has been shown to play a critical role in inflammation [66–72]. PARP1 is required for nuclear factor kappa-light-chain-enhancer of activated B (NF-kB) transcription and its activation [73]. It also increases mRNA stability of pro-inflammatory genes and enhances release of pro-inflammatory molecules such as high-mobility group box protein 1 (HMGB1) [74, 75]. MARylating PARPs also participate in modulating the immune response [76]. MARylating and non-enzymatic PARPs such as PARP9/11/12/13/14 have broad spectrum antiviral properties and their gene expression was shown to increase during inflammation [76].

Consistently, in the human placental microarray, we observed an increase in the transcript levels of such MARylating and non-enzymatic PARPs (PARP4, PARP9, PARP12 and PARP14) in the inflammation-driven PE placentas. While we did not observe an increase in the transcript levels of PARP1 or other PARylating enzymes in this dataset, we did observe increased PARylation in the human placentas from the inflammation-driven PE patients, in the trophoblasts treated with TNF-ɑ and in the LPS treated rat placentas. A number of studies also demonstrated activation of PARP1 mediated PARylation in inflammatory conditions with no reported increase in the PARP1 gene or protein levels, indicating that cellular stress or inflammation may induce increased PARP1 enzymatic activity without increasing its expression [75, 77, 78]. Apart from having an inflammatory signature, we also observed oxidative DNA damage in the human and rat inflammation-driven PE placentas. Both PARP1 and PARP2 participate in DNA damage repair [33]. Thus, it is likely that various PARylating and MARylating enzymes are responsible for the decline in placenta NAD^+^ levels during inflammation-driven PE. Due to the known substantial contribution of PARP1 in inflammation, its inhibition has been shown to reduce inflammation and restore cellular NAD^+^ levels in several studies [69–72]. However, PARP1 inhibition or knockout in pregnant rodent models has caused fetal defects suggesting its essential role during fetal development [79–82]. Several studies suggest that PARP1 is required for cell survival under mild oxidative stress as it participates in DNA damage repair [83–85]. Conversely, under conditions of severe oxidative stress, excessive activation of PARP1 has been shown to induce cell death due to energy exhaustion by consuming NAD^+^ [86]. As placental oxidative stress is observed in PE, it is likely that activation of PARP1 is required for the adaptation to the stress; and later its excessive consumption of NAD^+^ leads to pathological consequences. As such, boosting NAD^+^ levels present an effective and safe treatment option for inflammation-driven PE.

It is now well established that reduced NAD^+^ content is a hallmark of defective energy metabolism [87]. This is now recapitulated in the placenta, where our study suggests that chronic inflammation in the placenta leads to a decline in NAD^+^ availability and mitochondrial dysfunction (Fig 5). Importantly, we show that therapeutically targeting the NAD^+^ signalling pathways and sustaining NAD^+^ levels throughout pregnancy maintains mitochondrial function, reduces inflammation and oxidative damage, thus, preventing PE disease development (Fig 5). NAD^+^ boosting has been proven to be effective in various pro-inflammatory disease models [19, 21, 28, 36, 65, 88]. Moreover, NR has been shown to be orally bioavailable and safe for human consumption [27, 89–91]. Several non-pregnant human studies, including in healthy/obese/overweight individuals and in patients with Ataxia or Alzheimer’s or Parkinsons disease, have demonstrated that NR is effective in elevating NAD^+^ levels and can improve systemic metabolic function [27, 92–97]. NR was also shown to reduce circulating inflammatory markers in humans [91], suggesting that it could be particularly effective for use with inflammation-driven PE patients. As mentioned earlier, NAM, has been found to be effective in hypoxia-driven PE mouse models [37, 38] in preventing hypertension and fetal growth restriction by reducing glomerular endotheliosis and improving metabolic profile of fetal brain [37, 38]. That being said, there is certainly a strong argument to be made that boosting NAD^+^ levels may have the potential to combat against all subclasses of human PE [8], thus making it a very promising therapeutic intervention.

**Figure 5:**
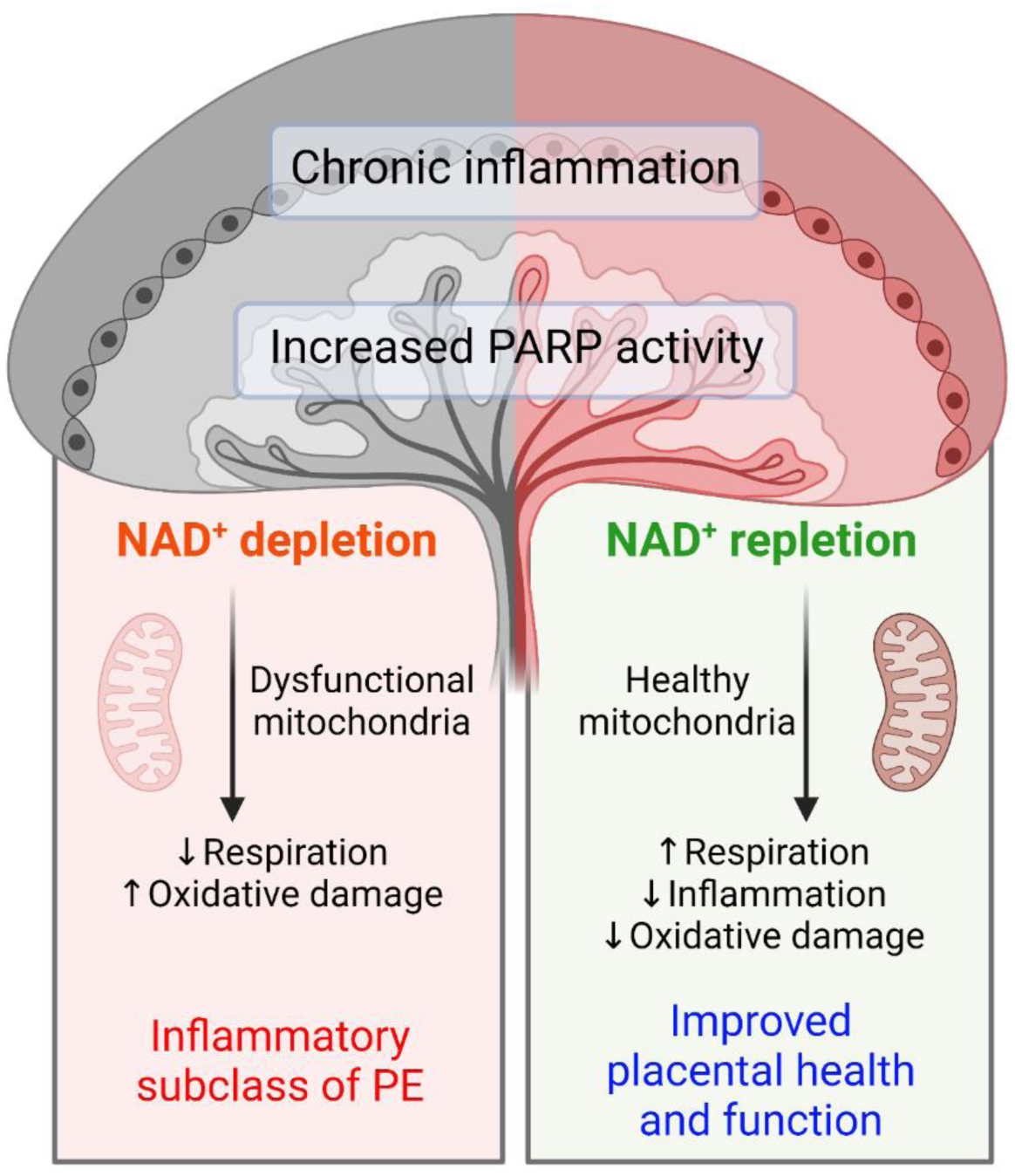
Effect of NAD^+^ boosting on inflammation-driven PE. Our findings suggest that a decline in placental NAD^+^ levels due to PARP hyperactivity in inflammatory PE leads to mitochondrial dysfunction and oxidative damage while NAD^+^ boosting strategies, through NR treatment, improves placental mitochondrial function and resolves placental inflammation and oxidative damage, thus, preventing the development of inflammatory PE (Created with BioRender.com on April 19^th^, 2023).

In conclusion, we showed for the first time that NAD^+^ plays a critical role in maintaining healthy placental function. We have mechanistically shown that depleted placental NAD^+^ level is a signature of human inflammatory PE and demonstrated that increasing NAD^+^ levels enhance placental function and thus prevents the development of the subclass PE3. Thus, in future, pre-clinical and clinical trials could be carried out to test the effectiveness of NAD^+^ boosting strategies, such as supplementing with vitamin B3 derivatives, during pregnancy or once PE has been diagnosed – to establish its treatment potential.

## Materials and Methods

### Study design

This study aimed to investigate the role of NAD^+^ in inflammation-driven PE (PE3), and to test the therapeutic potential of NAD^+^ boosting in an animal model of inflammation-driven PE. We hypothesized that there is a decline in placental NAD^+^ levels due to hyperactivity of NAD^+^ consuming enzymes and that it leads to impaired placental mitochondrial function, collectively contributing to the development of PE. Thus, replenishing NAD^+^ could be an attractive preventative strategy in this subclass of PE patients. We have used human placenta samples to profile placental NAD^+^ levels, activity of NAD^+^ consuming PARP enzymes and mitochondrial content/protein expression. We next performed cell culture experiments using SVneo/HTR8 trophoblast cell line to understand how NR can benefit placental cells under an inflammatory condition by treating the cells with TNF-α, a pro-inflammatory cytokine whose expression is increased in the placenta of inflammation-driven PE patients [3]. Finally, we employed an LPS-induced rat model of inflammation-driven PE to test the potential of NR in preventing PE development. In this model, we evaluated BP, fetal/placental weights, placental morphology, inflammation, mitochondrial function, and oxidative stress. We also assessed placental transcriptome and NAD^+^ metabolism to understand how boosting NAD^+^ may work to prevent PE development.

### Frozen Human placenta tissue

All human tissue experiments were carried out according to University of Ottawa and Mount Sinai Hospital research ethics board guidelines and approval (University of Ottawa REB protocol# H08-17-08, Mount Sinai Hospital REB protocol# 20-0037-E). For the measurement of placental NAD^+^ and mitochondrial content, a total of 128 frozen placenta samples and matched histological sections were purchased from the Research Centre for Women’s and Infants’ Health BioBank (RCWIH), Mount Sinai Hospital, Toronto, Canada. Samples included term and preterm placentas from healthy controls (n=43), individuals with chronic hypertension (n=23), gestational parental-driven PE subclass (PE1; n=16), hypoxia-driven PE subclass (PE2; n=42) and inflammation-driven PE subclass (PE3; n=17) patients. PE was defined as the onset of hypertension (systolic pressure > 140 mm Hg and/or diastolic pressure > 90 mm Hg) after 20 weeks of gestation coupled with evidence of maternal end-organ dysfunction [2]. Classification of the molecular subclass was done by microarray (confirmed by qPCR analysis) and histopathology analysis, as previously described [3, 98]. For comparison purpose, control healthy term and preterm placentas were combined as a singular healthy “control” group, since PE placentas included both term and preterm placentas.

Detailed methods of microarray gene expression analysis, protein PARylation and NAD(H) measurements, mitochondrial protein and oxidative stress measurements are provided in the appended supplement (Supplementary 5)

### Cell culture

The HTR-8/SVneo cell line, derived from human first-trimester placental villus explant, was purchased from ATCC (CRL-3271). Cells were cultured in Gibco® 1X RPMI-1640 (Life Technologies™) medium supplemented with 10% fetal bovine serum (FBS), at 37 °C, 5% CO_2_ and ~20% O_2_ condition. Methods of protein PARylation and NAD(H) measurements, mitochondrial respiration assay and OXPHOS protein expression; and trophoblast functional (invasion) assay are provided in the appended supplement (Supplementary 5). Briefly, cells were treated with TNF-ɑ (10mg/ml) to induce inflammation and NR (250 μM) was co-treated for specified timepoints to test its potential to improve trophoblast mitochondrial and cellular function.

### Animal study

All animal experiments were carried out according to University of Ottawa animal care ethics and guidelines (protocol# HS2923). Sprague Dawley rats (Charles River Laboratories International, Inc.) were kept in a room with 23°C temperature and a 12:12-hr light-dark cycle. Estrus cycle was confirmed by doing vaginal smears to allow timed mating [99]. Vaginal smears were checked the following day to identify presence of sperm to confirm pregnancy. Additionally, body weights were recorded daily to track pregnancy. The following experimental groups were included:

I. **LPS (n=10):** Lipopolysaccharide (Escherichia coli O55:B5, L2880-10MG, Sigma-Aldrich) was solubilized in sterile normal saline and animals were intraperitoneally (IP) injected from GD 13-18 at an daily incremental dose from 20ug-70ug/kg/day, as shown previously [100].
II. **NR (n=10):** NR was administered by oral gavage from GD 1-19. NR was solubilized in drinking water and sterile filtered and given at a concentration of 200 mg/kg/day.
III. **LPS + NR model (n=10):** NR and LPS was administered as mentioned above.
IV. **Saline (n=10):** Controls received oral gavage of sterile drinking water from GD 1-19 and saline IP injections from GD 13-18.

Animals were sacrificed on GD 19, fetal and placental weights and litter size were recorded and organs were collected for future analysis.

Detailed methods of pregnancy outcome, placental histomorphology, inflammation, gene expression, protein PARylation and metabolite measurements, placenta mitochondrial functionality assay and protein expression, and oxidative stress measurements are provided in the appended supplement (Supplementary 5).

## Statistical analysis

Statistical analysis was performed using GraphPad Prism software (Version 9.5). Error bars in graphical presentations indicate mean ± standard deviation (SD). Depending on the treatment groups, an unpaired t-test or one-way or two-way ANOVA with a Holm-Šídák’s multiple comparisons test was applied. For animal experimentations, robust regression, and outlier removal (ROUT) method (available on GraphPad Prism) was applied to exclude any outliers. A p value of < 0.05 was considered statistically significant.

## Author Contributions

Conceptualization, F.J., G.V, K.J.M and S.A.B.; data collection and analysis, F.J., G.V, Y.C., A.N-A., A.E.G; writing, F.J.; review and editing, F.J., K.J.M and S.A.B.; project supervision and funding, K.J.M and S.A.B. All authors have read and agreed to the published version of the manuscript.

## Funding

This research was funded by the Canadian Institutes of Health Research (CIHR) Project Grant #PJT-153055 to S.A.B. and K.J.M. F.J. is supported by Frederick Banting and Charles Best Canada Graduate Scholarship from CIHR (FRN-167027).

**Supplementary Figure 1:** Placental NAD^+^ related metabolite-NAM levels and mitochondrial content in human PE subclasses. A) Placental NAM levels were determined by LC/MS. B-C) Placental mitochondrial content was determined by measuring activity of two markers, citrate synthase and complex IV, by performing enzymatic activity assays (n=17-41 placentas/group). One-way ANOVA with Holm-Šídák’s multiple comparisons test. Error bar indicates standard deviation (SD). D) Heatmap of downregulated mitochondrial proteins in hypoxia-driven PE2 placentas compared to control placentas (n=5-6 per group). Shown are only proteins that met the False Discovery Rate (FDR) threshold of 0.05. Control= control term and preterm, CH = chronic hypertension control, PE 1 = Gestational parent-driven PE subclass, PE2 = Hypoxia-driven PE subclass, PE2 = Inflammation-driven PE subclass.

**Supplementary Figure 2:** NAD^+^ boosting in HTR8/SVneo cells under an inflammatory condition. Total cellular NAD^+^ quantification was performed on cells treated with 10ng/ml TNF-α and/or 250 μM NR for 24 hrs (n=3). One-tailed unpaired t-test, *P<0.05. Error bar indicates standard deviation (SD). TNF-α= Tumor necrosis factor, NR= nicotinamide riboside.

**Supplementary Figure 3:** Effect of NAD^+^ boosting on placental transcriptome of a rat model of inflammation-driven PE. Principal component analysis (PCA), Heatmap, Volcano plot and GO term analysis were performed between LPS and LPS+NR treated rat placentas (n=4-5 litter/treatment, 1placenta/litter).

**Supplementary Figure 4:** Effect of LPS and NAD^+^ boosting on placental expression of NAD^+^ consuming enzyme CD38, NAM levels and mitochondria. A-B) Placental expression of NAD^+^ consuming enzyme CD38 were determined by western blot with anti-CD38 antibody (n=5 litter/treatment, 1placenta/litter). C) NAM levels were determined by LC/MS (n=5 litter/treatment, 1placenta/litter), D) Mitochondrial complex-IV activity was measured by Oxygraph assay (n=4-8 litter/treatment, 6-8 placentas/litter). E-M) Placental OXPHOS, OPA1, CLPP and YME1L1 protein levels were determined by western blot (n=5 litter/treatment, 1placenta/litter). Two-way ANOVA with Holm-Šídák’s multiple comparisons test, *P<0.05,**P<0.01,***P<0.001. Error bar indicates standard deviation (SD).

