## Supplementary for "NAD^+^ depletion and altered mitochondrial function are key to the establishment of placental dysfunction in an inflammatory-driven subclass of preeclampsia"

#### Supplementary 1

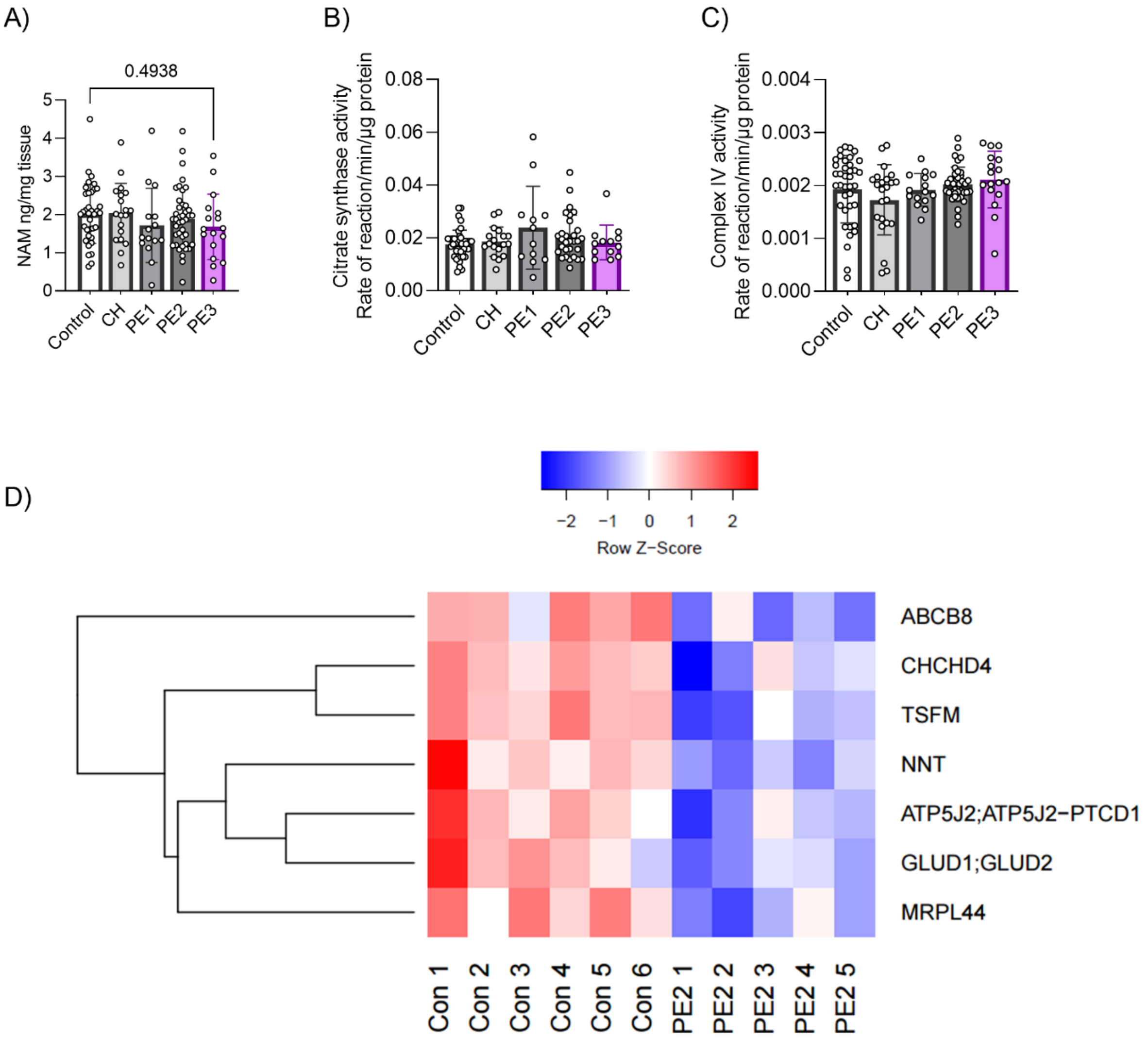

**Supplementary 2**

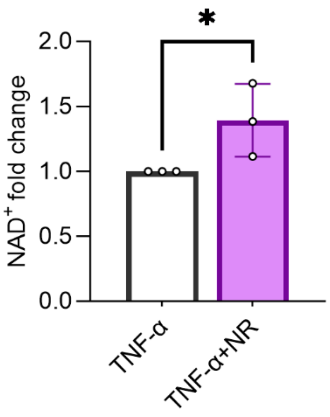

**Supplementary 3**

A)

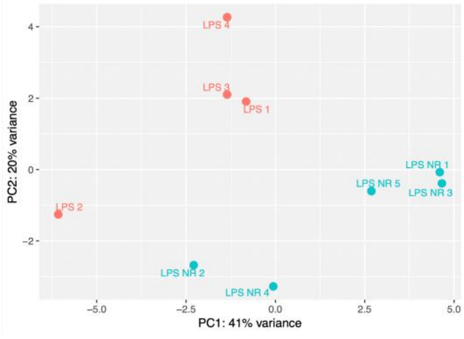


B)

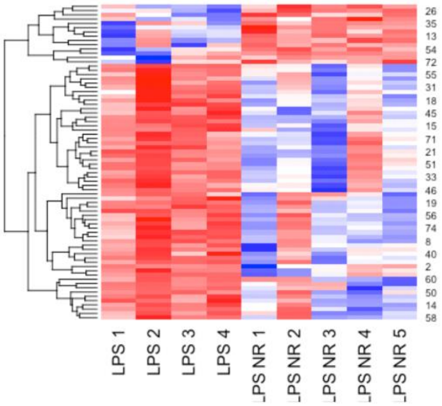


C)

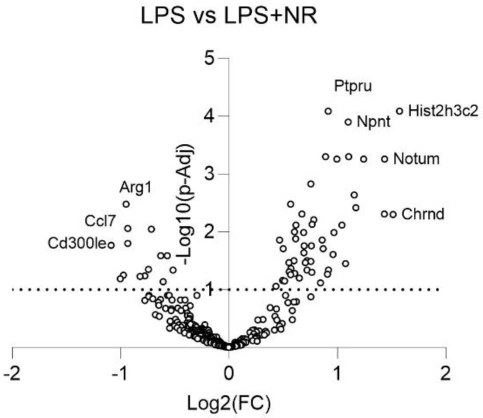


D)

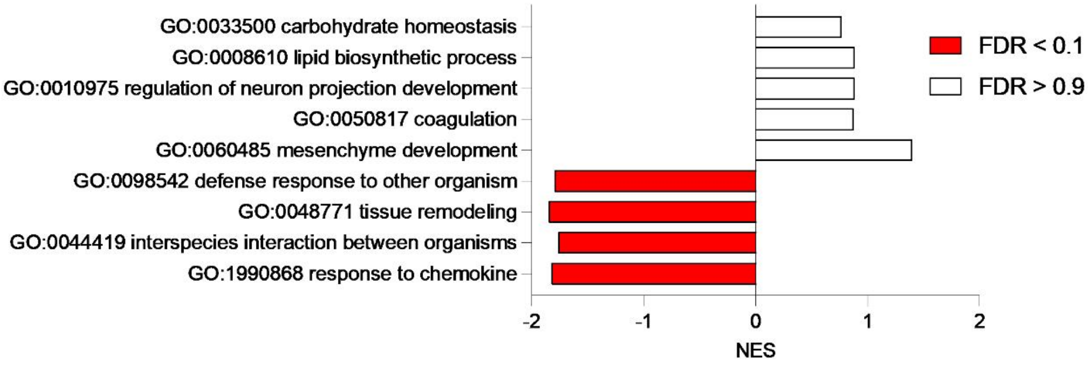

### Supplementary 4

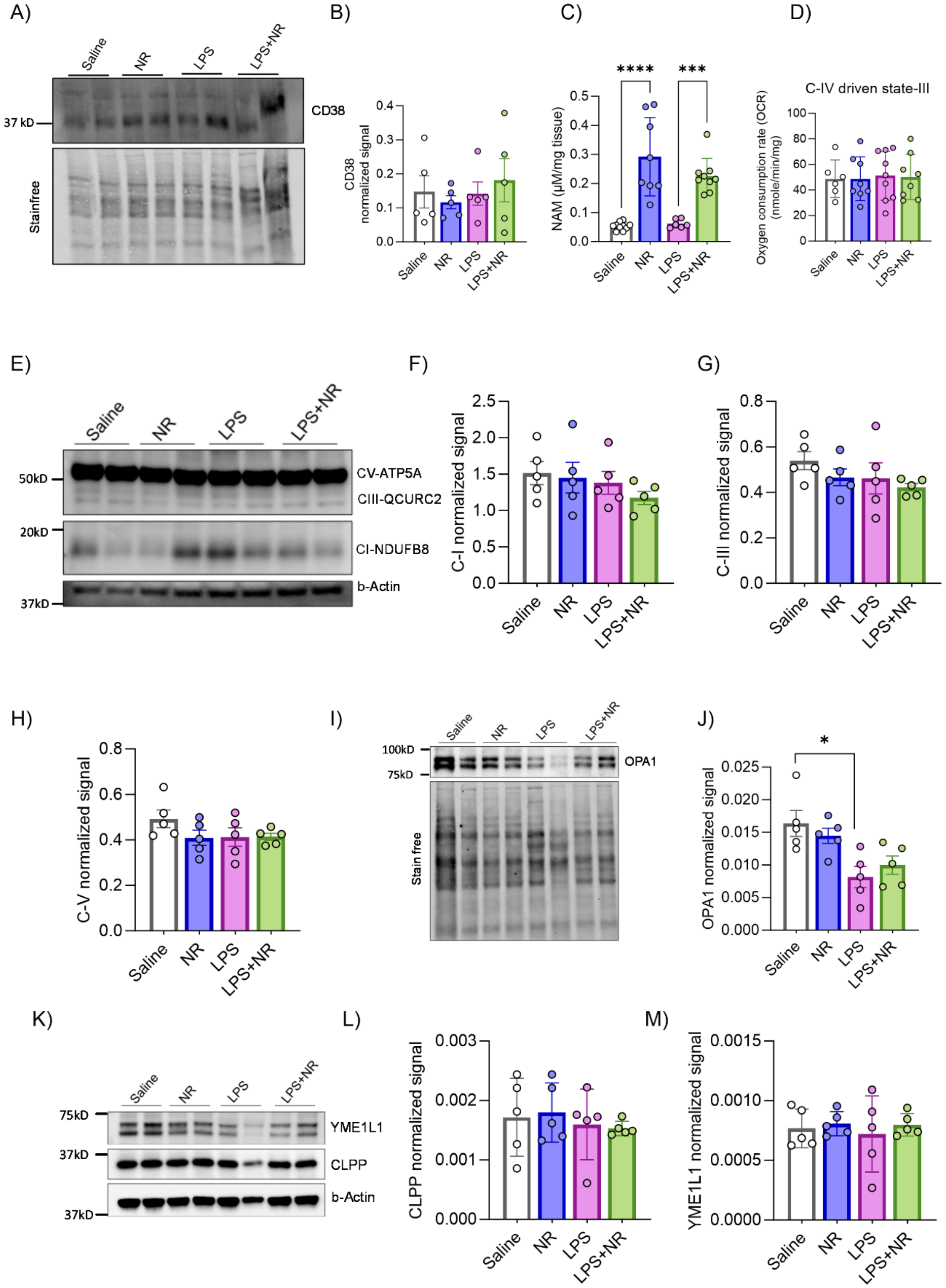

#### Supplementary 5

#### Materials and Methods

1. Human sample processing

*1.1 Microarray analysis of gene expression:*

A microarray dataset previously generated by our group and available at the Gene Expression Omnibus (GEO) database (GSE75010) was used to assess PE subclass-specific expression of inflammatory markers and $NAD^+$-consuming enzymes [3]. This microarray dataset was generated from frozen human placenta tissues that were purchased from RCWIH, Mount Sinai Hospital, Toronto, Canada. Briefly, RNA isolation was performed using Trizol and RNAeasy spin columns. Human Gene 1.0 ST Array chips (Affymetrix) was used to perform microarray at the Princess Margaret Genomics Centre (Toronto, Canada). The heatmap of expression of key genes of interest was generated in R studio.

*1.2 Measurement of PARylation*

Paraffin embedded placental tissue sections (5-µm), mounted on glass slides, were purchased from RCWIH, Mount Sinai Hospital, Toronto, Canada. These specimens were collected from the same cases as the frozen tissue biopsies described above (Table 1) and were used to assess protein PARylation and oxidative stress (section 1.5 below) within the placenta for each of the three PE subclasses and controls. Sections were dewaxed in xylene, and hydrated in a sequence of 100%, 95%, 80%, 70% and 50% ethanol, and water. Sections were subjected to antigen retrieval using citrate buffer (0.1 M, pH 6.0) in microwave for 7 minutes and endogenous peroxidase blocking using

3% $H_2O_2$ in PBS for 30 minutes. To prevent nonspecific binding, protein block solution (DAKO, X0909) was utilized for 1 hour. Subsequently, the tissue sections were incubated overnight at 4°C with Anti-PAR antibody (1:250, Cell signalling 83732) diluted in PBS containing 1% BSA, overnight at 4°C. Then, slides were incubated with Alexa Fluor 595 goat anti-rabbit secondary antibody (1:4000, Invitrogen A-11012) and kept in dark for 1h at room temperature. Tissue was counterstained with DAPI (Promega, P36931). 5 snapshots of random regions of interest (20X magnification) of the placental villi (trophoblast) were selected for fluorescence intensity quantification using ImageJ software.

*1.3 Measurement of NAD(H)*

Flash frozen human placental samples (same cases as describe above) were pulverized in liquid nitrogen to generate powdered tissue used for NAD(H) and mitochondrial protein analysis (section 1.4 below). For measurements of NAD(H), 13-30 mg of powdered placental tissue was weighed in a 1.5 ml microcentrifuge tube and 1000 µl of solvent A (40:40:20 acetonitrile:methanol:water with 0.1M formic acid) was added. The mixture was vortexed for 10 sec and kept on ice for 3 min. 87 µl of solvent B (15% $NH_4HCO_3$ in water) was added, vortexed and kept on dry ice for 20 min to neutralize.  The sample was centrifuged at 16,000 g for 15 min at 4 °C and supernatant was collected for LC-MS analysis [101].

NAD, NADH and NAM were determined using validated liquid chromatography mass spectrometry/mass spectrometry (LC/MS/MS). After sample extraction, three internal standards [IS, 13C5NAD, 13C 2,6,7 NAM and 6,7-Dimethyl-2,3-di(2-pyridyl)-quinoxaline (DMDPQ)] were added to all samples [blanks, standards, quality controls (QC) and tissue extracts]. After vortexing and centrifugation, the supernatant was transferred to auto-sampler vails for injection. The chromatographic separation was carried on an LC system (Accela, Thermo). The column was a Synergi Polar-RP (5 cm x 4.6mm, 2.5 µm) column (phenomenex). Separation was performed at 30 °C with a gradient elution of acetonitrile-0.1% (v/v) formic acid in water at a flow rate of 0.5ml/min. The auto-sampler was set at 4 °C and the injection volume was 10 ul. Mass spectral analyses were accomplished on a quadrupole tandem mass spectrometer (TSQ Quantum Access MAX, Thermo) with electrospray ionization in positive mode. Multiple-reaction-monitoring (MRM) mode was used to detect all compounds. MRM transitions were m/z 646 to 136, 666 to 649, 123 to 80, 669 to 136, 127 to 83, and 313 to 246 for NAD, NADH, NAM, 13C5NAD,13C 2,6,7 NAM, and DMDPQ, respectively. The calibration ranges were 200-20000 ng/ml, 100-10000 ng/ml, 5-500 ng/ml for NAD, NADH and NAM, respectively. The intra-day and inter-day precision were all between 80-120% and all extracted samples were stable at 4 degrees for 24 hours.

*1.4 Measurement of mitochondrial proteins*

For measurements of mitochondrial proteins, 12.5 mg of powdered placenta sample was added to 0.75 mL of cold lysis buffer (1% SDS in 50 mM triethylammonium bicarbonate (TEAB), Thermo Scientific Product No. 90114), and homogenized by passing 20 times through 21 G needle. Lysate was centrifuged at 20,000 x g for 10 minutes at 4C. Supernatant was separated, protein concentration was measured using BCA Protein Assay Kit (Thermo Scientific Product No. 23227), and 100 ug of sample was transferred into a new tube and adjusted to a final volume of 100 uL with 100 mM TEAB. 10 mM Reduction buffer (from freshly prepared 1M dithiothreitol in water) was added and kept at room temperature for 1 hour. 20 mM Alkylation buffer (from 1M iodoacetamide in water) was added and kept at room temperature for 45 min in the dark. Six volumes of pre-chilled protein precipitation buffer (50% acetone + 50% ethanol + 0.1% acetic acid) was added, vortexed and allowed to precipitate overnight at -20C. The following day, samples were centrifuged in a microfuge for 10 minutes at 10,000 x g. The supernatant was discarded carefully, and the remaining precipitation buffer was evaporated for 20 minutes at room temperature without over drying. 100µg of precipitated protein pellets was resuspended with 100µL of 100mM TEAB. Trypsin (Trypsin Gold, Promega) was used to digest the proteins (1:100 trypsin to protein by weight) and kept at 37°C for overnight. Peptide labeling was performed using TMT10plex Isobaric Label Reagent Set plus TMT11-131C (A34808, Thermo Scientific), following manufacturer's instructions.

Following reconstitution in 20uL 2% acetonitrile and 0.1% formic acid in water, 2uL was injected for analysis. The system used consisted of an Ultimate 3000 coupled with an Exploris 480 mass spectrometer (ThermoFisher Scientific, San Jose, CA) equipped with a Nanospray Flex interface operated in positive ion mode. The mobile phases consisted of 0.1%(v/v) FA in water as buffer A and 0.1% (v/v) FA in 80% acetonitrile as buffer B. The analysis was done on a column of 75µm × 200 mm, packed also in-house with reverse-phase Magic C18AQ resins (3µm; 120-Å pore size; Dr. Maisch GmbH, Ammerbuch, Germany).

Briefly, the sample was loaded on the column using 98% buffer A at a flow rate of 1µL/min for 10min. Then, a gradient from 5% to 35% buffer B was performed in 60 min at a flow rate of 300nL/min. The MS method was set up following the TMT MS2 template in Xcalibur 4.3.73.11. Specifically, it consists of one full MS scan from 350 to 1200 m/z, with a resolution of 120 000, defined at m/z 200m, followed by data-dependent MS/MS scan within 2 seconds, with dynamic exclusion of 1, exclusion duration of 45 s, and mass tolerance of 10 ppm, at an intensity threshold of 5E3. The isolation window was set to 0.7 Da, and the HCD collision energy was 36. The resolution of MS2 was set to 45,000, with the first mass as 110 to include all the TMT tags. To improve the mass accuracy, all the measurements in the Orbitrap mass analyzer were performed with internal recalibration ("Lock Mass" at 445.120025). On the Orbitrap, the charge state rejection function was enabled, with only 2~5 charges included. The raw data were processed and analyzed using MaxQuant (version 1.6.6.0). A target-decoy database search was carried out for *Rattus norvegicus* UniProtKB/Swiss-Prot and UniProtKB/TrEMBL protein sequence databases including commonly observed contaminants and their reversed sequences. A precursor ion mass tolerance of 0.5 Da was used to carry out this database search. TMT reporter intensities were corrected using the error correction factors provided with the kit.

*1.5 Measurement of oxidative stress*

Placental tissue sections (5 um) were deparaffinized and subjected to antigen retrieval as mentioned in 1.2. DNA was denatured by treating the slides with 2N HCl for 5 minutes at room temperature, followed by a neutralization step in 1M Tris-base for 5 minutes at room temperature.  Non-specific binding sites were blocked by incubating the sections in 10% normal goat serum in PBS, 1h at room temperature. Next, tissues were incubated with 8-oxo-dG antibody (1:250, R&D Systems 4354-MC-050) diluted in

PBS containing 0.1% BSA, overnight at 4°C. Slides were incubated with Alexa Fluor 488 goat anti-mouse secondary antibody (1:4000, Invitrogen A11029) in PBS containing 0.1% BSA in dark for 1h at room temperature. Tissue was counterstained with 7AAD (1:50, Invitrogen A1310) diluted in water for 30 minutes at room temperature in the dark. Quantification was performed as described in 1.2.

#### 2. Cell culture measurements

##### *2.1 Measurement of PARylation*

The HTR-8/SVneo cells were cultured in Gibco® 1X RPMI-1640 (Life Technologies™) medium supplemented with 10% fetal bovine serum (FBS), at 37 °C, 5% $CO_2$ and ~20% $O_2$ condition. 50,000 cells were plated on a 12 well-plate and allowed to grow overnight. The following day the cells were treated with TNF-α (10mg/ml), with and without NR (150 µM- 1 mM), for 24 hrs. A supplemented radioimmunoprecipitation (RIPA) buffer was prepared, which included Roche Complete Protease Inhibitor Cocktail (05892970001, Roche Canada, Basel, Switzerland), Roche PHOStop Phosphatase Inhibitor Cocktail (4906837001 Roche Canada, Basel, Switzerland), 5 mM of sodium butyrate, 100 µM of fresh tannic acid, and 1 µM of olaparib. Cells were then washed with ice-cold PBS and RIPA lysis buffer was added to each dish. Lysate was collected by scraping with a cell scraper (Thermo Scientific™). The lysate was vortexed and kept in a rotor for 30 min for slow mixing. The lysate was then centrifuged at 20,000 relative centrifugal force (r.c.f.) at 4°C for 15 minutes, and the supernatant was collected.

The protein concentration was determined using the DC protein assay (Bio-Rad Laboratories, Hercules, California, USA), with Bovine serum albumin (BSA) standards prepared in RIPA buffer. The resulting supernatant was then transferred to new tubes and stored in a -80°C freezer. The samples were prepared with 4x Laemmli Buffer (Bio-Rad Laboratories, Hercules, California, USA Cat# 1610737) augmented with 10% β-mercaptoethanol (Fisher Bioreagents, BP176-100) and boiled for 5 minutes.

Next, western blotting was performed using 8%, 10%, or 12% SDS-PAGE gels, depending on the size of the protein of interest. The TGX Stain-free FastCast Acrylamide Kit (Bio-Rad Laboratories, Hercules, California, USA Cat# 1610183) with 10% Ammonium persulfate (APS) was used to make the gel. Following SDS-PAGE, the Stain Free gels were activated and then the proteins were transferred to TransBlot Turbo Mini size 0.2µm nitrocellulose or PVDF membrane (Bio-Rad) using the Trans-Blot TurboTransfer System ((Bio-Rad Laboratories, Hercules, California, USA Cat# 1704150EDU). To detect the total protein, the blot was imaged again under stain-free blot setting with automatic exposure time. Then the blot was blocked for one hour in blocking buffer [5% w/v bovine serum albumin (Sigma, SKU A7906) in TBS-T buffer (50mM Tris-HCl, pH 7.6; 150mM NaCl; 0.1% Tween)] while rocking gently at room temperature. The membranes were then incubated with primary anti-PAR antibody (Millipore Sigma, AM80-100UG) diluted in TBS-T buffer overnight at 4°C with gentle rocking. After three washes each for 10 minutes with TBS-T, the membrane was incubated for one hour with the matching IgG, horseradish peroxidase (HRP)-conjugated secondary antibody, rocking at room temperature. After three 10 min washes in TBS-T, detection was carried out on a ChemiDoc system (Bio-Rad Laboratories, Hercules, California, USA) using 1 ml Clarity (Bio-Rad Laboratories, Hercules, California, USA) or Clarity Max (Bio-Rad Laboratories, Hercules, California,

USA) enhanced chemiluminescent solutions. Later, images were retrieved and the quantification of blots was performed using either ImageLab or FIJI software.

*2.2 Measurement of NAD(H)*

250,000 HTR-8/SVneo cells were plated on a 6 $cm^3$ plate and incubated overnight in serum starved RPMI 1640 medium (Gibco® 1X RPMI-1640, Life Technologies™) at 37 °C, 5% CO2 and ~20% $O_2$ condition. The following day, 10% FBS containing RPMI 1640 medium was added, and cells were treated with 10 ng/ml TNF-α, with and without NR 250 μM, and incubated for 24 hrs. NAD(H) levels were quantified using a Biovision $NAD^+$/NADH kit (K337) according to manufacturer's instruction.

*2.3 Measurement of mitochondrial respiration and protein content*

Mitochondrial respiration or oxygen consumption rate (OCR) was measured by performing XFe96 seahorse assay. 2,500 cell/well was plated in seahorse cell culture plate in 180 ul of RPMI 1640 media with 10% FBS and incubated at 37°C, 5% $CO_2$ for 24 hrs. 12 hours prior to running the assay cartridge was hydrated. 200 uL of XF Calibrant (pH 7.4) was added to each well of the calibration plate and the cartridge was placed on the machine and sealed. The cartridge and calibration plate were placed in a 37°C incubator (no $CO_2$). 100 mL of seahorse media was prepared freshly before starting the assay. Media constituents:  8.3 g/L DMEM powder (no H2CO3), 2 g/L D-glucose, 1 mM pyruvate, 0.3 g/L glutamine and made in 90ml of sterile molecular grade water. pH was set at 7.4 and the media was topped up to 100 ml with molecular grade water and filter sterilized. Cells were washed with seahorse media 2x. The seahorse plate was placed in the 37°C incubator (no $CO_2$) for 45 minutes to allow the plate to equilibrate. All seahorse drugs were prepared in seahorse media and loaded in the cartridge. Calibration was performed, followed by measurements of the basal respiration. Oligomycin (1 μM) was injected to inhibit mitochondrial complex V to record non-phosphorylating OCR. FCCP (0.5 μM), a mitochondrial uncoupler was injected to record maximal OCR. Antimycin A (1 μM) was injected to determine non-mitochondrial respiration to subtract from the recorded basal and maximal respiration. Monensin (200 μM) was injected to record glycolytic capacity of the cells by recording extracellular acidification rates. Results were normalized by measuring protein concentration of each well via Bradford assay. Briefly, seahorse media was removed from plate using a multichannel pipette and cells were washed with 100uL PBS. 30 uL of the lysis buffer (1mM Tris-HCL- pH 7.4, 1mM EDTA, 0.5% triton 100x) was added to each well using a multi-channel pipette and mixed gently, with 5 uL from each well mixed with 195 uL of 1:5 Bradford reagent (Biorad) for protein quantification. Absorbance was read at 595nm using a spectrophotometer. The measurement was compared with BSA protein standards to get relative quantification.

Mitochondrial OXPHOS protein expression was assessed by western blotting. Briefly, HTR-8/SVneo cells were cultured as per conditions described in 2.1 [TNF-α (10mg/ml) ± NR (250 μM) for 24 hrs]. Western blotting protocol was similar to that described in 2.1, using the OXPHOS antibody cocktail (ab110413).

*2.4 Measurement of invasive capacity*

250,000 HTR8/Svneo cells were plated on 6cm$^3$ plates. After overnight incubation, cells were treated with TNF-α (10ng/ml) ± NR (250μM) in 10% FBS containing RMPI1640 media for 48hrs. Media was changed at 24hrs, with replenishment of TNF-α (and/or NR treatment. Matrigel coated invasion inserts were used in a boyden chamber invasion assay. Briefly, a 24 well-plate containing Matrigel inserts were removed from freezer and warmed to room temperature (30 minutes). 750ul of serum free warm RPMI 1640 media was added to the bottom of each well, with an additional 500ul media added on top of each insert and incubated at 37° C for 2hrs to rehydrate the Matrigel. HTR8/Svneo cells were then trypsinized, neutralized, and transferred to top portion of the chamber, onto of the Matrigel (50,000 cells/well) in serum depleted RPMI1640 media. Serum enriched media was placed in the bottom of the chamber, promoting cellular migration through the Matrigel. The cells were incubated in the chamber overnight (with all the treatments) at 37° C, at 5% $CO_2$ and ~20% $O_2$ condition.

After a 24 hr incubation, media was removed from top and bottom of the chamber, with the insert removed and washed with PBS (2x). Then, 700uL of 10% Formalin was added to the wells, fixing the adherent cells to the insert at room temperature for 10 minutes. Inserts were removed from the fixative, and wash 2x with PBS. Inserts were then placed in 100% methanol for 20 min at room temperature to permeabilize the cells. Inserts were then washed 2x with PBS, and adherent cells stained with hematoxylin for 10 minutes at room temperature. Inserts were washed once more with PBS, followed by tap water twice. Using a cotton swab, any cells remaining in the top chamber (non-invaded cells) were removed. The insert was allowed to air dry, following which the membrane was cut out and mounted on a glass slide.

#### 3. Animal model measurements

##### *3.1 Measurements of pregnancy outcomes*

Maternal mean arterial blood pressure (MAP) was assessed across the last portion of pregnancy (GD14-19) for each of the four different treatment groups (control, NR alone, LPS alone, LPS + NR; n=7-8/group). Measurements were captured using the CODA® high-throughput non-invasive tail cuff measurement system (Kent Scientific Corporation). Animals were first acclimatized to the system for 4-5 days. Prior to recording, animals were stabilized for 5-10min, and body temperature was checked. Then BP was recorded for 20min on each testing day.

At GD19, animals were sacrificed. Litter size, number of resorptions, fetal weights and placental weights were collected. Placentas were collected and flash frozen for RNA, metabolite, and protein measurements. One placenta per litter were fixed in 4% PFA for histopathology.

*3.2 Measurement of placenta histomorphology*

PFA-fixed mid-line placenta tissues (n=1 placenta/litter; n= 10 litters/ group) were paraffin embedded, sectioned at 5 uM thickness and placed on glass slides. Sections were deparaffinized, rehydrated and stained with H&E using standard protocols [102]. Whole-slide image were captured using Axio Scan.Z1 microscope at 20X magnification. Measurements of total placenta mid-line cross-sectional area, as well as cross-sectional areas of each of the individual regions of the placenta (labyrinth zone, junctional zone and decidua) were analyzed using ZEN 3.1 blue edition.

*3.3 Measurement of placenta gene expression*

RNA was isolated from frozen placenta tissues (n=1 placenta/litter; n=10 litters/group) using standard Trizol® methods. Purified RNA was submitted to StemCore Laboratories at the Ottawa Hospital Research Institute for RNA-Sequencing as follows. RNA quantification and quality was assessed using a Qubit HS RNA assay (ThermoFisher Scientific, Q32852) and the Fragment Analyzer Standard Sensitivity RNA assay (Agilent, DNF-471-0500). All samples had an RNA Quality Number (RQN) above 8.0. DNA library was prepared using the Illumina Stranded mRNA Preparation Kit (Illumina). Libraries were quantified using the Qubit Double Stranded DNA HS kit (ThermoFisher Scientific, Q33230) and integrity confirmed with the High Sensitivity NGS assay on an AATI Fragment Analyzer (Agilent). DNA libraries were normalized, pooled, and diluted. 75-basepair single-end sequencing reads were obtained from pooled DNA libraries using the Illumina NS500 75 cycle high output kit on an Illumina NextSeq 500 Sequencer.

Sequencing results were separated by library barcoding, and barcodes removed using Illumina BaseSpace. Subsequently, data was imported into the Galaxy Platform maintained by GenAP (http://www.genap.ca) and hosted by the Digital Research Alliance of Canada. Quality of sequencing was assessed using the FastQC function. Sequences were aligned to the rat genome (mRatBN7.2) and read quantification was performed using the HISAT2 and feature Counts functions, respectively. Count normalization, differential gene expression, and principal component analysis between control and LPS treated group was performed using DESeq2 and an FDR of 10%. Using only the differentially expressed genes with LPS treatment, to determine genes rescued with NR treatment, a subsequent analysis using DESeq2 identified genes that changed between LPS and LPS with NR groups with an FDR of 10% was performed.

Z-scores were calculated from normalized counts of differentially expressed genes and heatmaps generated in R studio. Gene Set Enrichment Analysis was performed using WebGestalt (http://www.webgestalt.org/) and the Reactome and Gene Ontology Biological Process databases.

*3.4 Measurement of placenta TNF-α protein content*

Placenta TNF-α protein expression was measured using Quantikine® Rat TNF-α Immunoassay immunosorbent assay (R&D Systems, Minneapolis, MN). Frozen placenta tissue (n=1 placenta/litter; n=10 litters/group) was pounded in liquid $N_2$ and weight about 40mg for homogenising 5mM EDTA/PBS extraction buffer with protease inhibitor cocktail (complete™ Roche 11836170001). TNF-α ELISA was then carried out according to manufacturer's protocol.

*3.5 Measurement of protein PARylation*

Flash frozen placenta (n=1 placenta/litter; n=10 litters/group) was pulverized, and 15-25 mg of tissue was used. RIPA buffer was added to the tissue and homogenized using 21G syringe. The homogenate was vortexed for 30 min to ensure complete homogenization. The homogenate was centrifuged at 20,000 relative centrifugal force (rcf) for 15 minutes at 4°C, and the supernatant was transferred to a 1.5 mL microcentrifuge tube. Western blotting was performed as described in 2.1 using anti-PAR antibody (Millipore Sigma, AM80-100UG).

*3.6 Measurement of NAD(H)*

30-40mg of powdered placenta tissue (n=1 placenta/litter; n=10 litters/group) was added to 300ul of extraction buffer. To prepare a homogeneous solution, the solution was drawn up and down through a 21-gauge syringe needle. Quantification was performed using the Biovision $NAD^+$/NADH kit (K337), according to the manufacturer's instructions.

*3.7 Measurement of NAM and ADPR*

Placental ADPR and NAM levels were quantified at the metabolomics core facility of University of Ottawa. Approximately 40 mg (+/- 10%) of powdered placental tissue (generated from n=1 placenta/litter; n=10 litters/group) was taken and added to 1.8 ml pre-chilled (-80°C) 80% methanol. Four 2.8 mm ceramic beads were added to each tube and the samples were homogenized at 2,000 rpm for 45 seconds using a MagNa Lyser bead homogenizer for up to 3 cycles. To keep samples, cool between cycles, the tubes were kept on dry ice for 1 min. Following homogenization, 1.8 ml of pre-chilled dichloromethane (-20°C) and 900 µl of ice-cold molecular grade water were added to the homogenate. For lipid removal, the mixture was vortexed and kept on ice for 10 minutes for partitioning. Samples were then centrifuged at 4,000 rpm at 1°C for 10 min and the upper aqueous phase was carefully collected and dried at 4°C using a refrigerated centrifugal concentrator (Labconco Refrigerated CentriVap Benchtop Vacuum Concentrator).

For quantification, 0.5 ug/ml of deuterated d5-Trp (Tryptophan) and 0.05 ug/ml of d4-NAM were used as Internal Standard (IS) in 75% acetonitrile to reconstitute the study samples. The mix was further tested in samples to check carryover or matrix effect on the endogenous metabolite level. Samples were reconstituted to a final volume of 50 µL. Samples were used undiluted for ADPR measurements, and 10X diluted (in 75% acetonitrile) for NAM measurements. Mass spectral analyses were accomplished on a Quadrupole time-of-flight (QTOF) mass spectrometer (MS) (Agilent 6545B), with electrospray ionization in positive mode connected to an ultra-high performance liquid chromatography system (Agilent 1290 InfintyII). 2 μL of samples was injected into the system., and Agilent Q-TOF Quantitative Analysis software (Agilent technologies inc. Santa Clara, CA, USA) was utilized to process and analyze the data. The calibration curve was optimized for each metabolite. Following peak detection, peak area was integrated using the Agile2 integrator. A >10 signal/noise ratio and accuracy of 80%-120% was used. Metabolite concentrations were normalized by the IS signal and adjusted to account for dilution factors. Metabolite concentrations were then normalized by the tissue weight.

*3.8 Measurement of mitochondrial respiration and protein content*

Freshly isolated mitochondria were used to measure mitochondrial respiration rates using a high-resolution oxygraph assay. Fresh placenta tissue (n=6-8 placentas/litter; n=10 litters/group) was collected to isolate mitochondria according to a modified version of a previously published protocol [103]. Briefly, placentas were washed in ice-cold PBS and transferred to a beaker containing buffer A [300mM sucrose mM, 10mM Tris-HCl, 1mM EGTA-Tris Base and 0.1% bovine serum albumin (BSA, fatty acid-free); pH 7,2, kept on ice]. Placentas were cut into smaller pieces and washed with buffer A twice to eliminate residual blood. Placentas were minced and transferred to a Potter-Elvehjem homogeniser (30ml) and homogenised at 500rpm for 6 strokes. The homogenate was centrifuged for 10 min at 1,000xg at 4 °C. The supernatant was transferred to a clean tube and centrifuged again with the same settings. Next, the supernatant was collected using a syringe to eliminate fat layer. The supernatant was collected in a clean tube and centrifuged for 10 min at 8,000xg at 4 °C. The supernatant was removed, and the pellet was resuspended in 2ml of buffer B (300mM sucrose, 10mM Tris-HCl and 0.05mM EGTA-Tris Base; pH 7) and centrifuged for 10 min at 8,000xg at 4 °C. This step was repeated. 200µl of MiR05 respiration buffer (110 mM D-sucrose, 60 mM lactobionic acid, 20 mM taurine, 20 mM HEPES, 10 mM KH2PO4, 3 mM MgCl2, 0.5 mM EGTA, 1 g/L fatty-acid free BSA; pH 7.1 at 23 °C)[104, 105] was added to the pellet and gently mixed to make a mitochondrial suspension. Mitochondrial protein content was measured by performing detergent compatible colorimetric (DC) protein assay (Biorad).

Mitochondrial respiration rates were determined by using an Oxygraph assay, according to a published protocol [106]. Briefly, mitochondria (2 mg/ml in MiR05 respiration buffer) were added to the chamber and baseline traces were recorded.  Substrates and inhibitors of mitochondrial respiration were injected in the following series: i) 5mM glutamate and 2.5mM malate, a Complex I substrate; ii) 1mM ADP, to measure Complex-I driven state-III respiration  ; iii) 1mM amytal to inhibit Complex I; iv) 5mM succinate to measure Complex-II driven state-III respiration  ; v) 5µM antimycin A to inhibit Complex III to prevent electron transfer to complex IV. This allows to measure complex-IV mediated respiration by adding substrate exogenously; vi) 5/0.3 mM Tetramethyl-p-phenylenediamine/ascorbate (TMPD)/Ascorbate) to measure Complex-IV driven state-III respiration, and vii) Finally, Potassium cyanide (KCN) to inhibit complex IV.

Mitochondrial protein expression was measured in flash frozen placenta tissues (n=1 placenta/litter; n=10 litters/group). Samples were prepared as described in 3.5 and western blotting was performed according to methods described in 2.1, using anti-

OXPHOS (Abcam, ab110413), Anti-OPA1 antibody (Abcam, ab42364), YME1L1 (Proteintech, 11510-1-AP) and anti-CLPP (Abcam, ab124822) antibodies.

*3.9 Measurement of placenta oxidative stress.*

Measurement of placental p-H2A.X protein expression, a marker of double stranded DNA-breaks and oxidative stress, were assessed using frozen placenta tissues (n=1 placenta/litter; n=10 litters/group) by western blot. Samples were prepared as described in 3.4 and western blotting was performed according to 2.1. using p-H2A.X antibody (Cell Signaling #2577). In parallel, paraffin embedded placenta tissue sections from the same litters (n=1 placenta/litter; n=5 litters/group) underwent deparaffinized, antigen retrieval and blocking for non-specific binding as described in section 1.5. Tissue sections were incubated overnight at 4°C with pH2A.X antibody (1:300, Cell Signaling #2577) diluted in PBS containing 1% BSA. Slides were then incubated with secondary antibody for 1 hour at room temperature, followed by incubation with streptavidin-HRP for 30 minutes, using the LSAB2 system-HRP (DAKO, K0675). The reaction was developed using 3,3′-Diaminobenzidine (DAB) (Sigma Aldrich D5637) plus $H_2O_2$. The tissues were counterstained with Harris hematoxylin and mounted in entellan. Whole slide images were captured using a Zeiss Axio Scan Z.1 scanner at 20X magnification and analyzed using QuPath software [107]. The decidua, junctional zone, and labyrinth were delimited to perform layer-specific automated positive cell counting. The results were expressed as pH2A.X positive cells per $mm^2$
